# Anti-α4β7 therapy targets lymphoid aggregates in the gastrointestinal tract of HIV-1 infected individuals

**DOI:** 10.1101/346684

**Authors:** Mathieu Uzzan, Minami Tokuyama, Adam K. Rosenstein, Costin Tomescu, Ivo N. SahBandar, Huaibin M. Ko, Louise Leyre, Anupa Chokola, Emma Kaplan-Lewis, Gabriela Rodriguez, Akihiro Seki, Michael J. Corley, Judith Aberg, Annalena La Porte, Eun-young Park, Hideki Ueno, Ioannis Oikonomou, Itai Doron, Iliyan D. Iliev, Benjamin K. Chen, Jennifer Lui, Timothy W. Schacker, Glaucia C. Furtado, Sergio A. Lira, Jean-Frederic Colombel, Amir Horowitz, Jean K. Lim, Nicolas Chomont, Luis J. Montaner, Lishomwa C. Ndhlovu, Saurabh Mehandru

## Abstract

Herein, we present the first human study of anti-α4β7 therapy in a cohort of HIV-1 infected subjects with mild inflammatory bowel disease. α4β7^+^ gut homing CD4^+^ T cells are early viral targets and contribute to HIV-1 pathogenesis, likely by seeding the gastrointestinal (GI) tract with HIV. Although, simianized anti-α4β7 monoclonal antibodies (Mab) have shown promise in preventing or attenuating the disease course of SIV in Non-Human Primate studies, the mechanisms of drug action remain elusive and the impact on HIV-1 persistence remains unanswered. By sampling the immune inductive and effector sites of the GI tract, we have discovered that anti-α4β7 therapy led to a significant and unexpected attenuation of lymphoid aggregates, most notably in the terminal ileum. Given that lymphoid aggregates serve as important sanctuary sites for establishing and maintaining viral reservoirs, their attrition by anti-α4β7 therapy has important implications for HIV-1 therapeutics and eradication efforts, and defines a rational basis for the continued evaluation of anti-α4β7 therapy in HIV-1 infection.

**One Sentence Summary:** Anti-α4β7 integrin therapy results in attrition of lymphoid aggregates within the gastrointestinal tract of HIV-1 infected individuals

## INTRODUCTION

Lentiviruses such as human immunodeficiency virus (HIV) and simian immunodeficiency virus (SIV) are uniquely adapted to infect activated, memory CD4^+^ T cells that are specifically enriched at mucosal surfaces (*1*). Consequently, mucosal tissues including those of the gastrointestinal (GI) tract play a critical role in disease pathogenesis during acute (*2, 3*) and chronic HIV-1 infection (*4*).

The GI tract can be immunologically sub-classified into inductive and effector sites (*5*). Peyer’s patches (PP) and isolated lymphoid follicles (ILF), intrinsic to the bowel wall and mesenteric lymph nodes (MLN), extrinsic to the bowel wall serve as the major immune inductive sites. Naïve T and B cells express integrin α4β7 (α4β7), that mediates their migration into the inductive sites through specific interactions with Mucosal Addressing Cell Associated Molecule-1 (MAdCAM-1) (*6*). Notably, the expression of α4β7 on naïve T and B cells is significantly lower than its expression on memory cells (*6*). PP-resident Dendritic cells (DC) prime naïve T and B cells and simultaneously induce the expression of high levels of α4β7 integrin in a retinoic acid and TGF-β dependent fashion (*7*). These α4β7^hi^, gut-primed, antigen-experienced memory cells egress into the draining lymph and subsequently into circulation and home to immune effector sites such as intestinal lamina propria, again via specific interactions between MAdCAM-1 and integrin α4β7 (*8*).

The pathogenesis of HIV-1 infection intersects with intestinal homing pathways at multiple levels that are yet poorly understood. GI-resident CD4^+^ T cells are preferentially targeted during acute HIV and SIV. Regardless of the route of infection and mode of virus delivery, intestinal CD4^+^ T cells are profoundly depleted during the earliest stages of HIV-1 and SIV infection (*9*). This strongly suggests that HIV-1, either cell free or cell-associated has evolved specific mechanisms to localize to GI tract during acute infection in order to infect CCR5-expressing (*10*) physiologically-activated memory T cells (*11, 12*) that are exceptionally HIV-1 susceptible (*2*). In this regard, studies have reported a direct interaction between α4β7 and the viral envelope (*13-15*). Thus, HIV-1 susceptible α4β7^+^CD4^+^ T cells may serve as Trojan horses, delivering the virus into the gut tissues.

Multiple lines of evidence demonstrate that α4β7-expressing cells represent early targets for the virus (*16-20*). This was highlighted in a recent report demonstrating that pre-infection frequencies of α4β7 on circulating CD4^+^ T cells may predict the risk of HIV-1 acquisition and disease progression independent of other T cell phenotypes and genital inflammation in a large cohort of at-risk South African women (*21*). Supporting this finding, STDs that have been linked with increased risk of HIV-1 acquisition, increase the frequency of α4β7^+^CD4^+^ memory T cells in both the mucosa and blood (*22, 23*).

Due to the critical role of α4β7^+^CD4^+^ T cells in viral pathogenesis, anti-α4β7 therapy has been considered in the management of HIV-1 infection. However, no human studies are available to date. In non-human primate (NHP) models, using simianized anti-α4β7 antibodies have shown promising results. Salient among these studies is the demonstration of disease prevention or an attenuated disease course when anti-α4β7 antibodies preceded low-dose repeated intravaginal SIV challenge (*24*). Additionally, a recent report found that SIV-infected macaques that were treated during acute infection with combination antiretroviral therapy (cART) and anti-α4β7 therapy (or isotype control) achieved long-term viremic control following treatment interruption (*25*). Despite multiple NHP studies, clear mechanisms underlying the potential efficacy of anti-α4β7 therapy in HIV (SIV) infection remain elusive.

While no HIV-related studies have been reported, anti-α4β7 therapy (Vedolizumab) has become a frontline strategy in the management of patients with inflammatory bowel diseases (IBD) (*26, 27*) where it has demonstrated strong efficacy and an outstanding safety profile (*28*). In order to determine its role in HIV-1 infection, we have studied a cohort of IBD patients with concomitant HIV-1 infection (*29*). Here we provide the report in humans on the safety, and the immunological and virological effects of anti-α4β7 therapy in HIV-1 infected patients receiving Vedolizumab (VDZ) therapy over 30 weeks, with detailed analyses in the GI tissue and in peripheral blood. The data reported herein generate novel insights into the mechanisms of action of anti-α4β7 therapy and provide a rational basis for its continued use in HIV-1 infection.

## RESULTS

### Vedolizumab was administered safely and without any Serious Adverse Events to patients with HIV-1 infection

Six patients (5 males and 1 female) with a median age of 51.7 years (interquartile range [36.8 - 62.2]) were followed prospectively for 30 weeks post-VDZ treatment. Five were receiving cART for a minimum of 5.6 years and had an undetectable viral plasma viral load at VDZ start (threshold of 20 copies/ml). One patient (583-016) was cART-treated for 9 months and had a plasma viral load of 156 copies/ml at the time of VDZ initiation. Detailed HIV characteristics are displayed in Table 1.

**Table 1.**
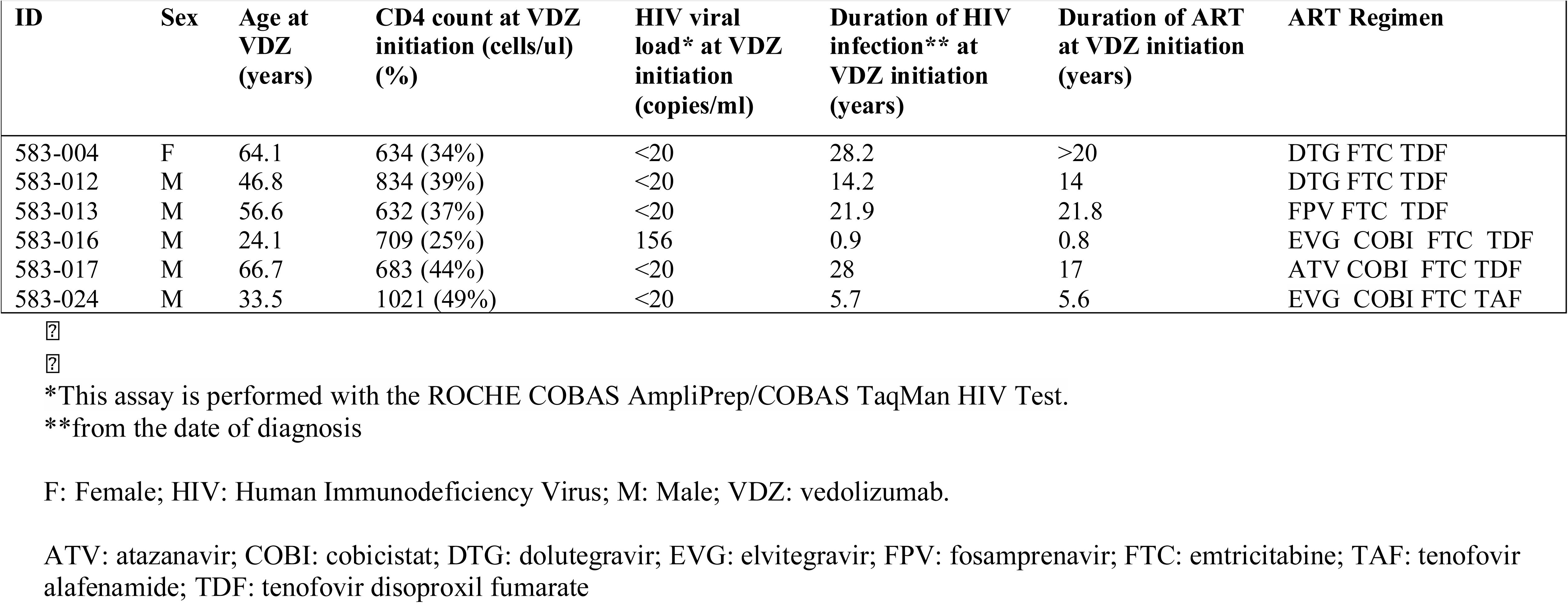
HIV-related clinical characteristics.

| ID | Sex | Age at VDZ (years) | CD4 count at VDZ initiation (cells/ul) (%) | HIV viral load* at VDZ initiation (copies/ml) | Duration of HIV infection** at VDZ initiation (years) | Duration of ART at VDZ initiation (years) | ART Regimen |
| --- | --- | --- | --- | --- | --- | --- | --- |
| 583-004 | F | 64.1 | 634 (34%) | <20 | 28.2 | >20 | DTG FTC TDF |
| 583-012 | M | 46.8 | 834 (39%) | <20 | 14.2 | 14 | DTG FTC TDF |
| 583-013 | M | 56.6 | 632 (37%) | <20 | 21.9 | 21.8 | FPV FTC TDF |
| 583-016 | M | 24.1 | 709 (25%) | 156 | 0.9 | 0.8 | EVG COBI FTC TDF |
| 583-017 | M | 66.7 | 683 (44%) | <20 | 28 | 17 | ATV COBI FTC TDF |
| 583-024 | M | 33.5 | 1021 (49%) | <20 | 5.7 | 5.6 | EVG COBI FTC TAF |

\*This assay is performed with the ROCHE COBAS AmpliPrep/COBAS TaqMan HIV Test.
\*\*from the date of diagnosis
F: Female; HIV: Human Immunodeficiency Virus; M: Male; VDZ: vedolizumab.
ATV: atazanavir; COBI: cobicistat; DTG: dolutegravir; EVG: elvitegravir; FPV: fosamprenavir; FTC: emtricitabine; TAF: tenofovir alafenamide; TDF: tenofovir disoproxil fumarate

All patients included in the study had very mild IBD activity (Table 2), characterized by mild proctitis and endoscopically normal appearing colonic and ileal mucosa with the exception of subject 583-016 where endoscopic inflammation was observed up to 25 cm from the anal verge. However, proximal parts of the colon and the TI, where study related biopsies were obtained, were normal in 583-016. Importantly, none of the study subjects had pancolitis, history of bowel surgery or previous use of biologic medications, all signifying severe disease (*30, 31*). Additionally, and as further evidence of mild IBD, 5/6 patients had normal levels of C reactive protein (CRP) at the time of starting VDZ (583-016 had a CRP of 16.5 mg/l at the time of recruitment). Finally, histology, arguably the gold standard for assessment of mucosal inflammation, showed only a mild increase in inflammatory cells limited to the rectum of 5/6 patients. Significantly, the terminal ileum (TI) and left colon (LC), sites where immunological and virological analyses were performed, were histologically normal in each of the study subjects (Suppl. Figure 1).

**Table 2.** IBD-related clinical characteristics.

| IBD characteristics |  |  |  |  |  |  |
| --- | --- | --- | --- | --- | --- | --- |
| ID | IBD type | Involved GI tract segments | Duration of IBD at VDZ initiation (years) | Age at IBD diag. (years) | Concomitant IBD therapies | CRP (mg/l) before VDZ initiation |
| 583-004 | UC | Recto-sigmoid | 5.2 | 58.9 | 5-ASA | 1.6 |
| 583-012 | CD | Right colon, Rectum | 1.3 | 45.5 | None | 1.4 |
| 583-013 | CD | Recto-sigmoid with history of rectal abscess and perianal fistula | 14.9 | 41.7 | 5-ASA | 2.3 |
| 583-016 | UC | Recto-sigmoid | 1 | 23.1 | 5-ASA | 16.5 |
| 583-017 | CD | Rectum with history of para-rectal abscess | 10 | 56.6 | None | 2.4 |
| 583-024 | UC | Recto-sigmoid | 12 | 21.5 | 5-ASA | 0.2 |
| Pre and post VDZ clinical and endoscopic assessments |  |  |  |  |  |  |
| ID | Clinical assessment pre VDZ |  | Endoscopic findings pre VDZ | Clinical assessment post VDZ | Endoscopic findings post VDZ |  |
| 583-004 | - 3-4 bm / 24 hours<br>- 50% with blood |  | Mild proctitis with aphthoid ulcerations | - 1-2 normal bm<br>- Non bloody | Improved mild proctitis |  |
| 583-012 | - 6-8 bm<br>- 25% with blood |  | Mild right sided colitis | - 2 bm<br>- Non bloody | No inflammation |  |
| 583-013 | - 4-5 bm<br>- Non bloody |  | Mild proctitis | - 1-2 normal bm<br>- Non bloody | Improved mild proctitis |  |
| 583-016 | - 6 bm<br>- 100% with blood |  | Ulcerated proto-sigmoiditis up to 25 cm from the anal verge | - 2-3 normal bm<br>- Non bloody | Healed inflammation |  |
| 583-017 | - 2 semi formed bm<br>- Non bloody |  | No inflammation | - 2 semi formed bm<br>- Non bloody | No inflammation |  |
| 583-024 | - 4-5 bm<br>- 25% with blood |  | Mild proctitis | - 1-2 normal bm<br>- Non bloody | No inflammation |  |

bm: bowel movements; CD: Crohn's disease; GI: Gastrointestinal; IBD: Inflammatory bowel disease; UC: Ulcerative colitis; VDZ: Vedolizumab; 5-ASA: 5-aminosalicylic acid

Patients were monitored with serial laboratory and clinical assessments. One of the 6 patients reported mild, self-limited nasopharyngitis, a previously reported adverse effect of VDZ (*32*). Two of 6 patients reported mild, self-limited headache and one patient had IV infiltration during infusion. No other adverse events (AEs) or serious adverse events (SAEs) related to VDZ were reported during the course of 30 weeks of follow up, highlighting the safety of this drug in our cohort of HIV-1 infected subjects. These data mirror previous, extensive data in IBD (*32*). Additionally, although outside the scope of the present report, the 6 patients continue to be on long term follow up while on VDZ therapy (∼ 1 year of follow up) and no AEs or SAEs have been observed to date, again emphasizing the safety of VDZ within our cohort of HIV-1 infected subjects.

### Anti-α4β7 therapy results in a significant reduction in B cell subsets within the GI tract

We defined a flow cytometric strategy to identify the known B cell subsets and plasma cells in intestinal mucosa and in circulation, identifying plasma cells as live CD45^+^CD38^hi^CD27^+^ cells and non-plasma B cells as live CD45^+^CD38^−^CD19^+^ cells. Among non-plasma B cells, naïve B cells were defined as CD45^+^CD38^−^CD19^+^IgD^+^IgM^+^ cells, while switched memory (SM) B cells were defined as CD45^+^CD38^−^CD19^+^IgD^−^IgM^−^ cells (Suppl. Figure 2).

The TI contains more lymphoid aggregates and therefore more non-plasma cell B cells while the LC harbors more lamina propria lymphocytes and therefore, mostly differentiated plasma cells (*5*). There was a clear dichotomy in the effects of anti-α4β7 therapy between the TI and LC, reflecting distinct cellular composition of these two intestinal sites. For example, we observed a striking decrease in non-plasma cell B cells (CD19^+^CD38^−^) in the TI of all 5 patients by flow cytometry. In contrast, in the LC which contains significantly fewer non-plasma B cells, the decrease was less pronounced (Figure 1 A and B). Among B cell subsets, both naïve and switched memory B cells were reduced in the TI post-therapy. Again, in the LC with significantly fewer total B cells, decrease in B cell subsets was less pronounced. (Figure 1C and D). Among GI plasma cells, no decrease was noted in either the TI or LC (Figure 1A and B).

**Fig. 1.**
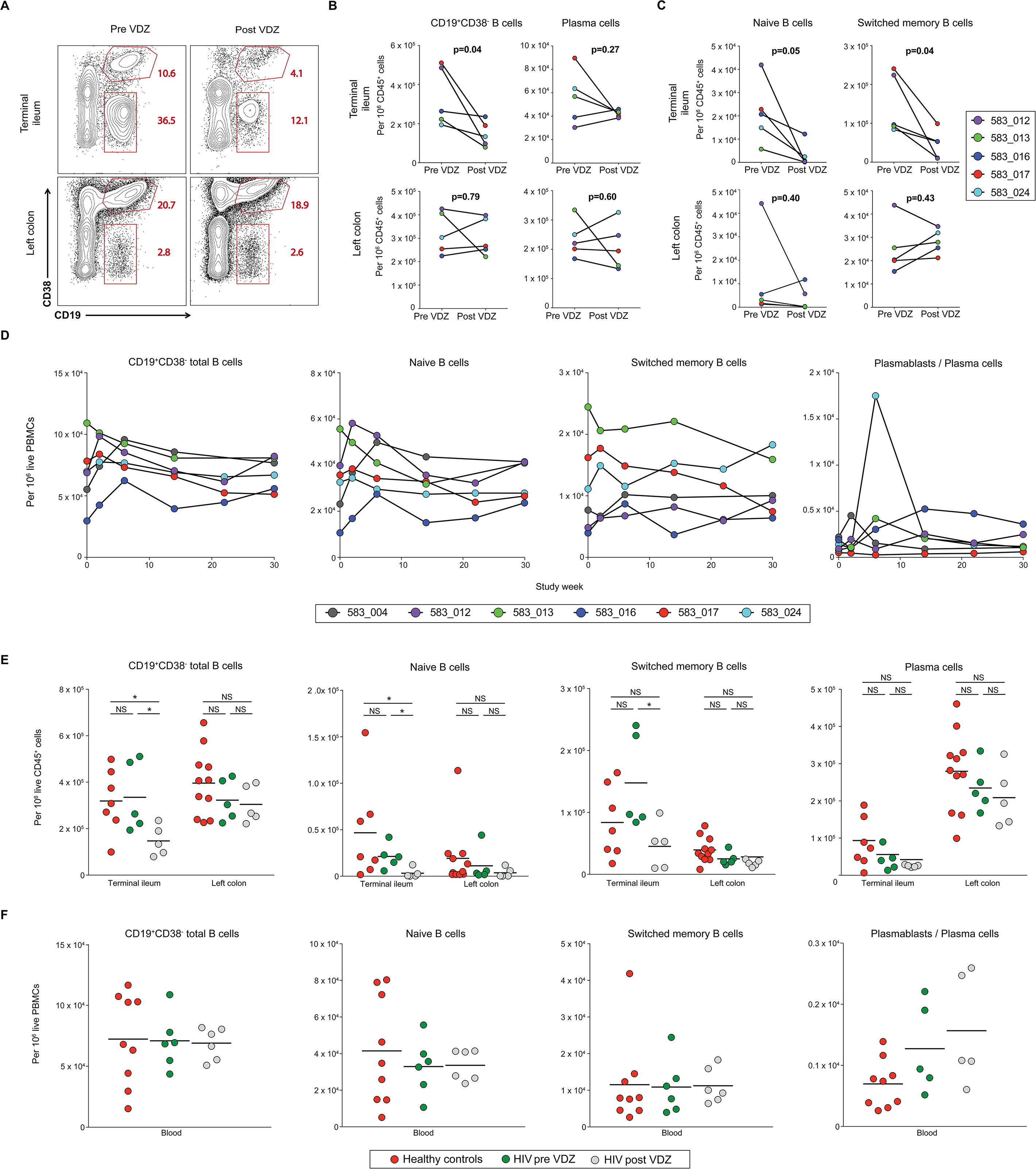
Anti-α4β7 therapy decreases the frequency of B cell subsets in the terminal ileum. (A-C) Change in the frequency of B cells and plasma cells in the GI tract post VDZ therapy. (A) Representative flow cytometry plots showing the expression of CD19 and CD38 among live, CD45^+^ cells derived from the terminal ileum (TI, top panels) and left colon (LC, bottom panels) of subject 583-017, before and at wk 30 post-VDZ. (B) Plots comparing the change in frequency of CD19^+^CD38^−^ total B cells and plasma cells (CD45^+^CD3 8^++^CD27^+^) in the TI (top) and LC (bottom) between pre- and post-VDZ treatment. (C) Plots demonstrating changes in the frequency of naive (CD45^+^CD19^+^CD10^−^CD38^−^IgM^+^IgD^+^) and switched memory B cells (CD45^+^CD19^+^CD10^−^CD38^−^IgM^−^IgD^−^) in TI (top) and LC (bottom). (D) Change in the frequency of total B cells, B cell subsets and plasma cells in the blood post VDZ therapy. In panels A-D each of the patients is represented with a unique color code (E) Group comparisons in the frequency of B cell subsets and plasma cells within the TI, the LC (F) Group comparisons in the frequency of B cell subsets and plasma cells in the peripheral blood. In panels E and F, Healthy volunteers are shown in red, HIV-IBD subjects pre-VDZ in green and HIV-IBD subjects post-VDZ in grey. Statistical values as indicated; *p<0.05

In circulation, while there was inter-individual variability, we observed a trend towards an early increase in all non-plasma B cells as well as in circulating plasmablasts (defined as Ki67^+^CD38^+^CD27^+^IgD^−^CD19^+/int^ cells, Suppl. figure 2) at week 2 post-VDZ initiation. These changes equilibrated over the course of 30 weeks of VDZ treatment (Figure 1D).

Next, we compared B cell composition between a cohort of healthy volunteers and VDZ treated patients. Compared to healthy volunteers, a significant decrease was noted in total CD19^+^CD38^−^ B cells in the TI post-VDZ treatment (Figure 1E). This was reflected in a significantly reduced frequency of naïve B cells, while changes in switched memory B cells and plasma cells in the TI were not significantly different between normal volunteers and VDZ-treated patients. In contrast to the TI, we did not observe significant changes in total B cells, B cell subsets (naïve and switched memory B cells), or plasma cells in the LC post-VDZ treatment, when compared to healthy volunteers (Figure 1E). Finally, no significant changes in circulating B cell subsets were noted post-VDZ treatment when compared to healthy volunteers, although circulating plasma cells showed a trend towards increase (Figure 1F).

Overall, all 5 patients presented a major decrease in non-plasma B cells (including both naive and memory subsets) in the TI with anti-α4β7 therapy.

### Anti-α4β7 therapy results in attrition of lymphoid aggregates within the GI tract

Next, we quantified GI B cells by immunohistochemistry (IHC) to confirm and further define the anatomical compartments showing changes in B cells post-VDZ. CD20^+^ non-plasma B cells were quantified per unit area in lymphoid aggregates and lamina propria in the TI and LC. Lymphoid aggregates were noted pre-treatment in 4/5 subjects in the TI and 2/5 subjects in LC. Remarkably, in the TI, there was a pronounced CD20^+^B cell reduction in TI-associated lymphoid aggregates post VDZ in all 4/5 subjects where lymphoid aggregates were detectable in both pre and post treatment biopsies. In the LC, marked reduction in lymphoid aggregate-associated B cells was noted in 1 subject while in the other, the reduction was more modest (Figure 2A and B). Expectedly, the frequency of lamina propria B cells was significantly lower than those in lymphoid aggregates. VDZ-treatment induced a reduction in lamina propria B cells in the TI and had a variable effect on lamina propria B cells in the LC (Figure 2A and 2B, Suppl. Figure 3).

**Fig 2.**
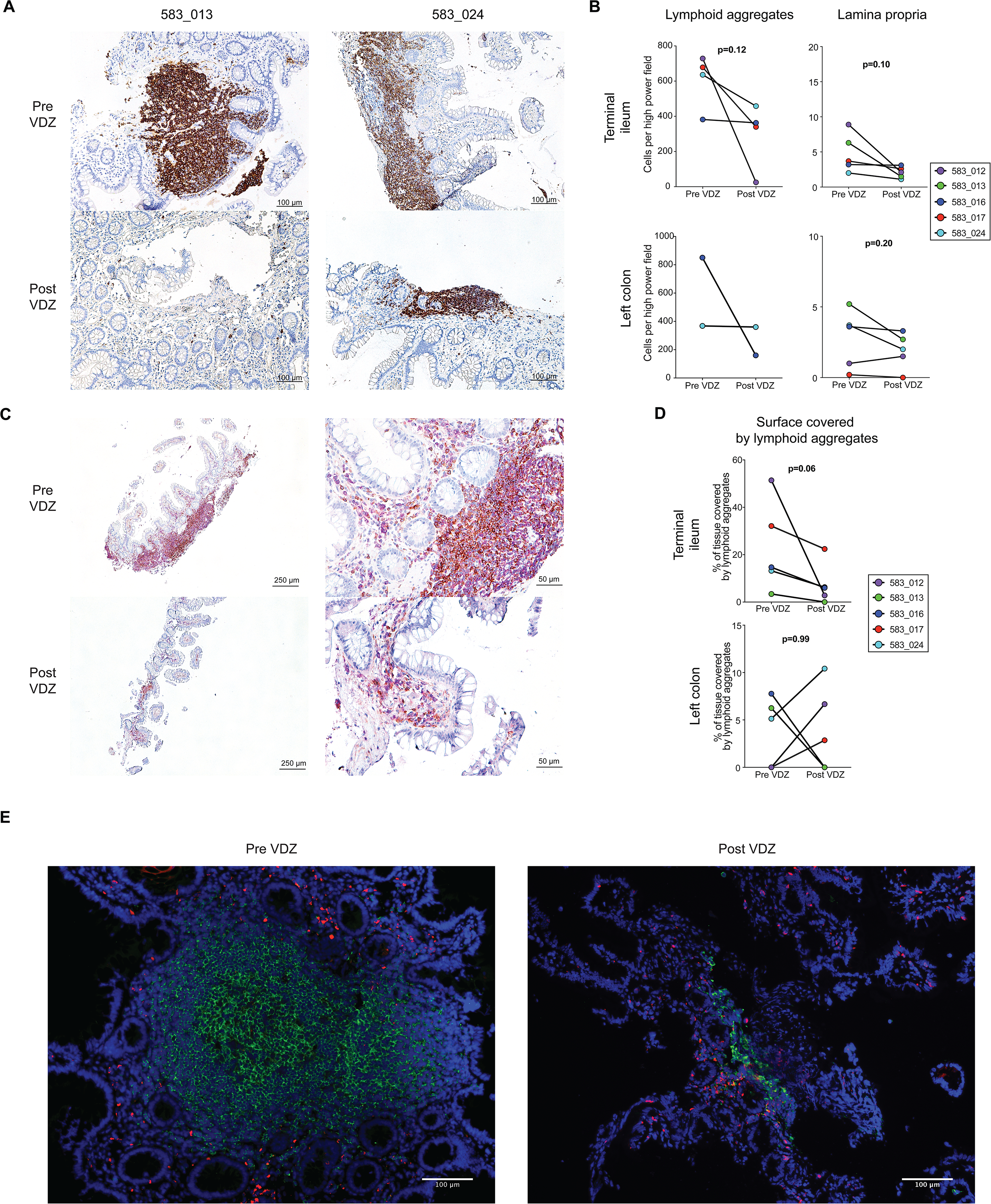
Anti-α4β7 therapy results in a significant attenuation of lymphoid aggregates, most pronounced in the terminal ileum. (A) Representative x10 magnification images of TI-derived biopsies immunostained for CD20 expression (brown) in 2 subjects (583-013 and 583-024) pre- (top panels) and post-VDZ (B) Quantitative analyses of the change in CD20^+^ B cells in the TI (top) and LC (bottom). Cell frequency was determined separately in lymphoid aggregates (left) and in lamina propria (right). (C) Representative images from subject 583-017 showing dual immunohistochemical staining with CD19 (pink) and CD4 (brown) pre- (top) and post-VDZ (bottom). Original magnification= 4X (left panel) and 20X (right panel) (D) Change in the percentage of tissue covered by lymphoid aggregates in the TI (top) and LC (bottom) in each of the subjects (E) Representative immunofluorescent image showing the expression of CD3 (red), CD20 (green) and DAPI (blue) from the TI of subject 583-024, pre-(left panel) and post-VDZ (right panel). Original magnification = 10X. In panels B and D each of the patients is represented with a unique color code. Statistical values are as indicated.

**Fig 3.**
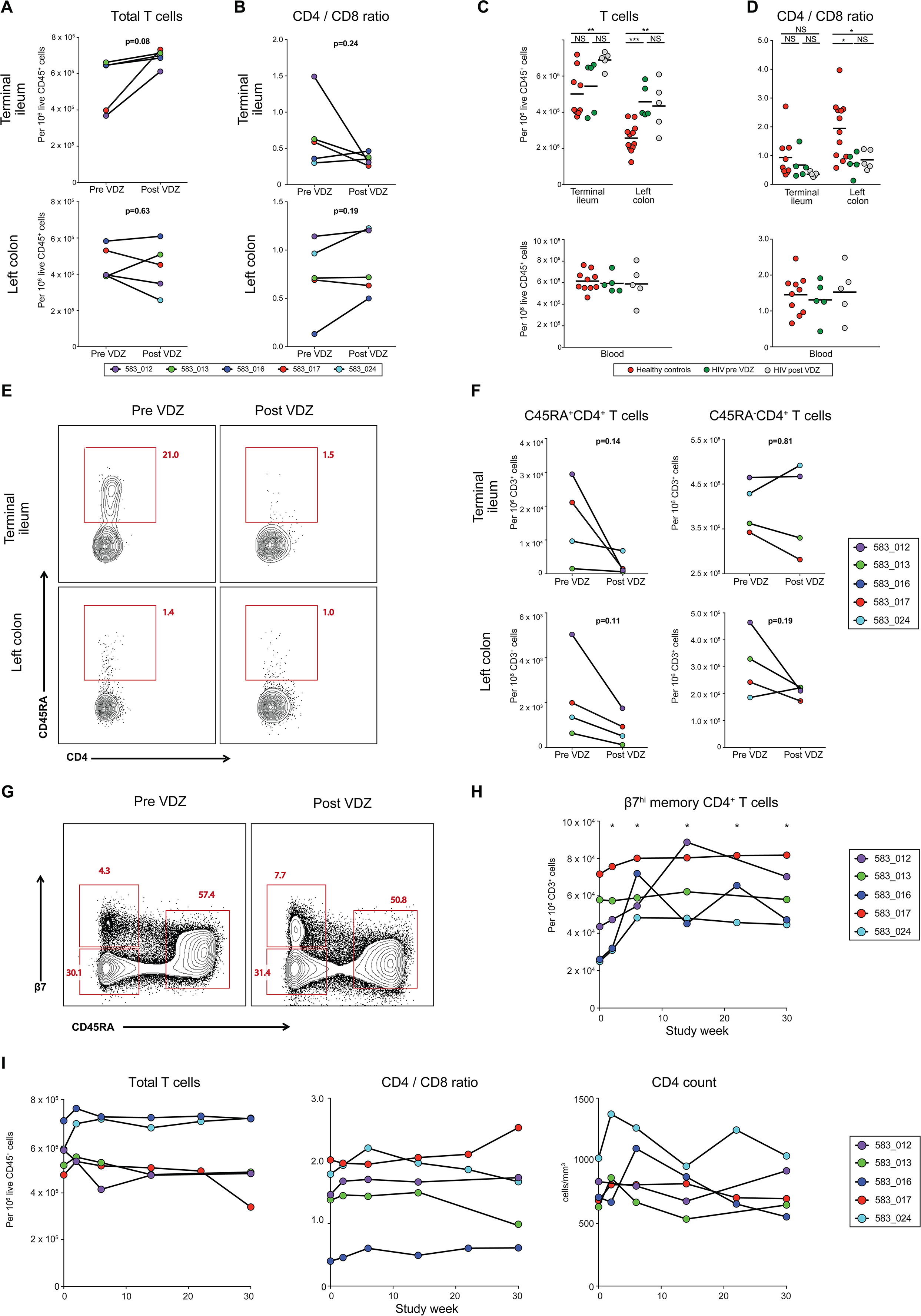
Anti-α4β7 therapy results in a decrease in naive CD4^+^ T cells in the terminal ileum. (A-B) Change in the frequency of T cells in the GI tract post VDZ therapy. (A) Change in the frequency of total CD3^+^T cells; and (B) CD4: CD8 T cells ratio in the TI (top panels) and LC (bottom panels) post-VDZ. Each of the patients is represented with a unique color code, statistical values are indicated. (C-D) Group comparisons between the frequency of T cells (C) and CD4:CD8 ratio (D) in the GI tract (top panels) and peripheral blood (bottom panels). Healthy volunteers are shown in red, HIV-IBD subjects pre-VDZ in green and HIV-IBD subjects post-VDZ in grey. *p<0.05, ** p<0.005, NS=not significant. (E-F) Change in the frequency of naïve and memory CD4^+^ T cells in the GI tract post-VDZ (E) Representative flow cytometry plots showing the expression of CD45RA on CD45^+^CD3^+^CD4^+^ T cells derived from the TI (top panels) and LC (bottom panels) of subject 583 −024 pre- and at wk 26 post-VDZ (F) Cumulative data showing changes in CD45RA^+^ (left) and CD45RA^−^ CD4^+^ T cell subsets within the TI (top panels) and LC (bottom panels). Notably, RA^+^ staining on CD4^+^ T cells was available on 4/5 patients that are shown here. (G-I) Change in the frequency of T subsets in the peripheral blood post-VDZ (G) Representative flow cytometry plots from subject 583-017 comparing the expression of β7 integrin and CD45RA on circulating CD4^+^ T cells. Pre-VDZ and at wk30 post-VDZ Three distinct populations are defined: β7 integrin^hi^CD45RA^−^, β7 integrin^−^ CD45RA^−^ and β7 integrin^int^CD45RA^+^. (H) Cumulative data showing changes in circulating β7 integrin^hi^CD45RA^−^ CD4+ T cells during VDZ therapy for each patient. In panels F-I, each of the patients is represented with a unique color code, statistical values are indicated. *p<0.05

Although the putative mechanism of action of anti-α4β7 therapy is to prevent the entry of α4β7^hi^ memory T cells into the intestinal lamina propria, to date, the published reports show no change in the frequency of lamina propria CD4^+^ T cells post-VDZ, either in SIV-infected macaques (*33*) or in humans with IBD (*34*). Importantly, the effects of VDZ on lymphoid aggregates, where cellular entry is also α4β7-MAdCAM dependent (*6*) remain unappreciated.

Having observed a significant decrease in non-plasma B cells in the TI, we hypothesized that anti-α4β7 therapy has a pronounced affect on lymphoid aggregates. To define lymphoid aggregates within tissue, we used IHC and quantified the surface area covered by lymphoid aggregates in each of the tissue sections pre- and post-VDZ (pathologists were blinded to the identity of the samples). In every subject, we observed a decrease in the percentage of total tissue surface covered by lymphoid aggregates post-VDZ in the TI (from 24.11 ± 19.3 % in average to 4.1 ± 2.9 %, Figure 2C and 2D, Suppl. Figure 3). Again, more variability was observed in the LC (from 3.8 ± 3.6 % on average to 3.9 ± 4.5 %), likely because lymphoid aggregates are less pronounced in the LC compared to the TI (*5*). In order to validate the IHC data, IF microscopy was performed to examine B and T cell populations in the lymphoid aggregates vs the lamina propria. Consistent with previous findings, we found a significant decrease in lymphoid aggregates post-VDZ (Figure 2E).

### Anti-α4β7 therapy results in a decrease in naive CD4^+^ T cells in the terminal ileum

SIV-macaque studies have demonstrated that anti-α4β7 therapy in combination with early cART enables better reconstitution of CD4^+^ T cells in the colonic mucosa compared to cART alone (*25*). We therefore investigated the impact of VDZ on T cell subsets, including CD4^+^ and CD8^+^ T cells in the GI tract and in circulation. In the TI, there was a trend towards an increased frequency of total CD3^+^ T cells post-VDZ treatment while CD3^+^ T changes in the LC post-VDZ were variable (figure 3A). Among T cell subsets, there was no significant change in the CD4:CD8 ratio in the TI or the LC (Figure 3B). When compared to healthy volunteers, there was a significant increase in CD3^+^ T cells in the TI and the LC post-VDZ (Figure 3C, top panel). In contrast to total T cells, there was no significant change in the CD4:CD8 ratios in the TI post-VDZ when compared to healthy volunteers. HIV-1 infected subjects had significantly reduced CD4:CD8 ratio in the LC when compared to healthy volunteers, consistent with published literature (*35*), and no significant VDZ treatment-related effects could be discerned in the CD4:CD8 ratio in the LC (Figure 3D, top panel). Finally, among circulating T cells, there were no significant differences in the total T cells or CD4:CD8 ratios between healthy volunteers or the study subjects, pre- or post-VDZ treatment (Figure 3C and 3D, bottom panels).

Among CD4^+^ T cell subsets, there were no significant changes in the memory CD4^+^ T cells (CD3^+^CD4^+^CD45RA^−^) post-VDZ in the TI or LC (Figure 3E and 3F). In contrast, we noticed a trend towards a decrease in naïve CD4^+^ T cells (CD3^+^CD4^+^CD45RA^+^) in the TI and LC post-VDZ (Figure 3E and 3F). Importantly, naïve T cells reside within the lymphoid aggregates while memory T cells are largely distributed in the intestinal lamina propria (*5*), these data again demonstrate a previously unappreciated effect of anti-α4β7 therapy on lymphoid aggregates in the GI tract.

In circulation, there was a significant rise in β7^hi^CD45RA^−^CD4^+^ T cells, shown to be uniquely susceptible to HIV-1 infection (*21*) at week 2, sustained over the duration of therapy (figure 3G and 3H). In contrast, while there was a short-term rise in total CD3^+^ T cells, CD4:CD8 ratios and absolute CD4+ T cell counts at week 2 post-treatment, over the course of 30 weeks, these counts tended to return to baseline (Figure 3I).

### Anti-α4β7 therapy results in a decrease in immune activation of CD4^+^ T cells in the terminal

Cellular activation, causally related to susceptibility to HIV-1 infection and HIV-1 latency (*36*), an independent marker of disease progression (*37, 38*), is significantly greater in the GI tract than the peripheral blood of patients with HIV-1 infection (*39*). In order to determine the impact of anti- α4β7 therapy on immunological activation in the GI tissue and peripheral blood, we assessed for the expression of CD38 on CD4^+^ and CD8^+^ T cells. A significant reduction in the frequency of CD4^+^CD38^+^ T cells was seen in the TI in all 5 subjects post-VDZ treatment (Figure 4A and 4B). In the LC, CD4^+^CD38^+^ cells decreased in 4 out of 5 subjects. Among CD8^+^ T cells, changes in CD8^+^CD38^+^ cells were not significant in the TI or the LC (Figure 4A and 4B). In the peripheral blood, while individual subjects showed a reduction in CD4^+^CD38^+^ cells and CD8^+^CD3 8^+^ cells, there was variability as a group and the results did not reach significance (Figure 4C). When compared with healthy volunteers, there was a trend towards reduced CD4^+^CD38^+^ cells in the TI, post-VDZ treatment. In contrast, among circulating T cells, the frequency of CD4^+^CD38^+^ and CD8^+^CD38^+^ cells was remarkably similar between healthy volunteers and VDZ-treated subjects (Figure 4D).

**Fig. 4.**
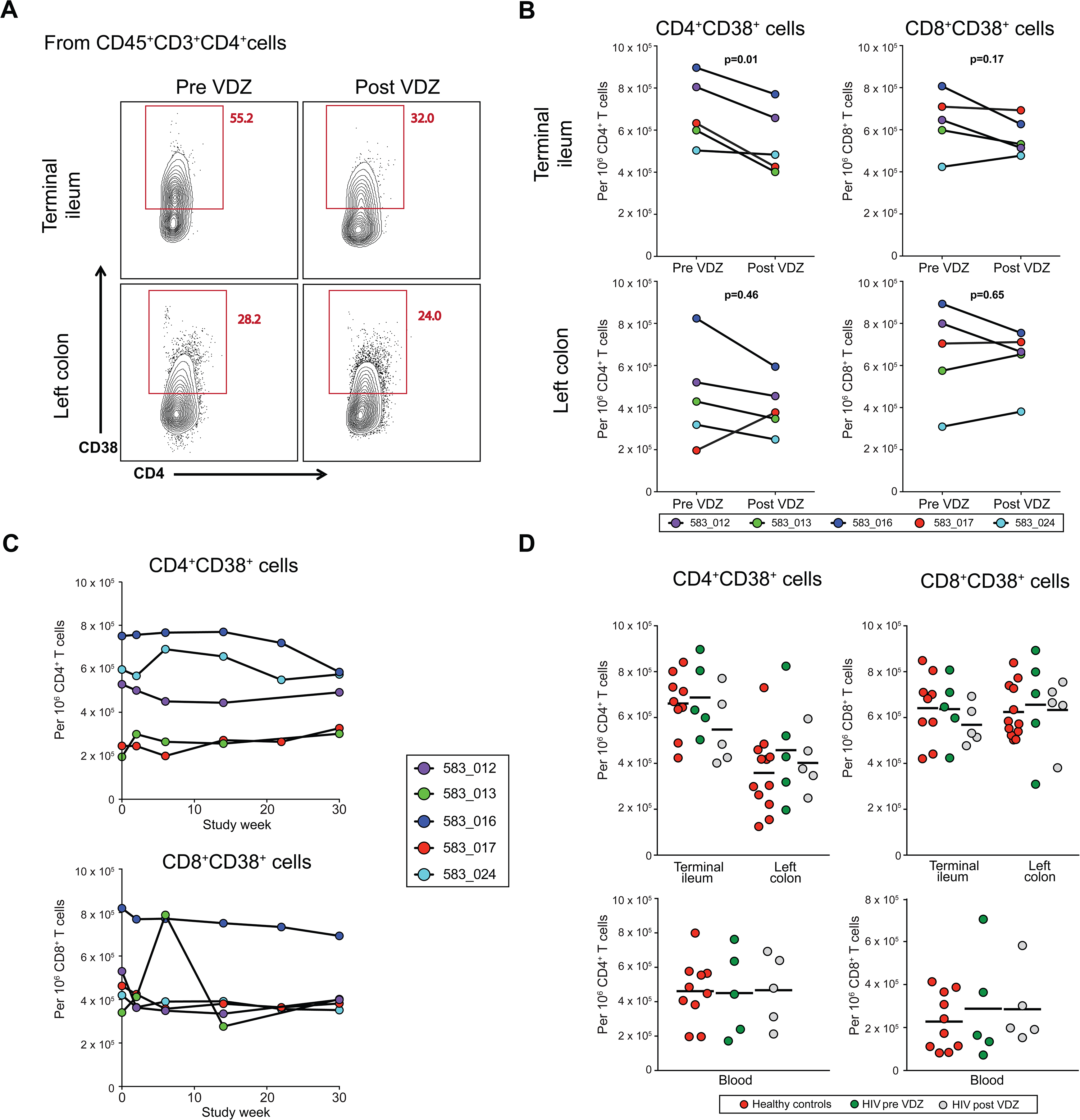
Anti-α4β7 therapy results in a decrease in immune activation of CD4^+^ T cells in the terminal ileum. (A-B) Change in the frequency of activated T cells in the GI tract post-VDZ (A) Representative flow cytometry plots showing the expression of CD38 on live CD45^+^CD3^+^CD4^+^ T cells derived from TI (top) and LC live cells in subject 583-013 pre-VDZ and at wk 26 post VDZ. (B) Cumulative data showing changes in the level of CD4^+^CD38^+^ T cells (left) and CD8^+^CD38^+^ T cells (right) in the TI (top panels) and LC (bottom panels) in each of the study subject postVDZ (C) Change in the frequency of CD4^+^CD38^+^ T cells (top) and of CD8^+^CD38^+^ T cells (bottom) in the peripheral blood post-VDZ In panels B and C, each of the patients is represented with a unique color code (D) Group comparisons between the frequency of CD4^+^CD38^+^ T cells (top left) and CD8^+^CD38^+^ cells (top right) in the GI tract and in the peripheral blood (botttom panels). Healthy volunteers are shown in red, HIV-IBD subjects pre-VDZ in green and HIV-IBD subjects postVDZ in grey.

### Anti-α4β7 therapy results in early changes in NK cell composition and activation that equilibrates over time

NK cells play a critical role in viral immunity including HIV with multiple lines of evidence supporting a role for both cytotoxic and regulatory functions in HIV-1 infection (*40*). Therefore, we analyzed the composition and the activation of NK cells at baseline and at wk 2 and wk 30 post-VDZ. We defined NK cells in three distinct subsets as follows: “cytolytic NK cells” were defined as CD56^dim^CD16^high^ NK cells, “cytokine secreting” NK cells as CD56^bright^CD16^low^ NK cells and CD56^null^ cells as CD56^low^CD16^high^ NK cells (Figure 5A). Strikingly, we found that in all patients the frequency of circulating CD56^bright^CD16^low^ “cytokine secreting” NK cells decreased at week 2 and increased back to a level similar to baseline at week 30 (Figure 5C). No clear trend was seen in the two other subsets (Figure 5B and 5D).

**Fig. 5.**
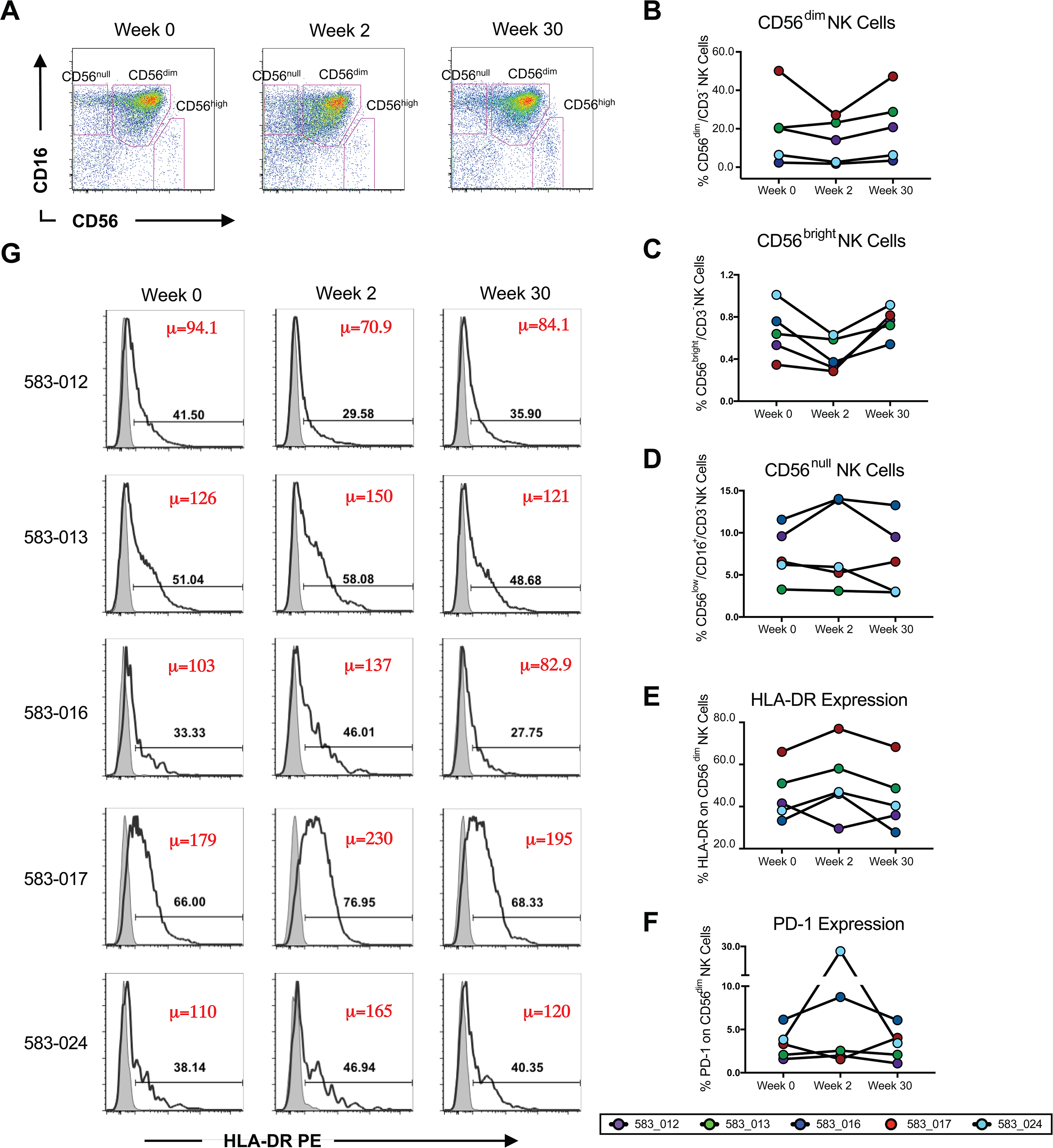
Anti-α4β7 therapy results in the activation of circulating NK cell subsets. (A) Changes in NK phenotype at weeks 0, 2 and 30 of therapy with Vedolizumab (VDZ). After exclusion of dead cells, monocytes, B cells and T cells; “Cytolytic NK cells” were gated as CD56^dim^CD16^high^ NK cells, “Cytokine secreting” NK cells were gated as CD56 ^high^ CD16^low^ NK cells and CD56^null^ cells were gated as CD56^low^CD16^high^ NK cells. (B-D) Composite graphs representing frequency of “Cytolytic” (B), “cytokine producing” (C) and “CD56^null^” NK cells (D) from each of the 5 subjects (color coded) at weeks 0, 2 and 30 of therapy with VDZ. (E-F) Individual evolution of the expression of the HLA-DR (E) and PD-1 (F) on “cytolytic” CD56^dim^CD16^high^ NK cells from 5 subjects (color coded) at weeks 0, 2 and 30 of therapy with Vedolizumab. (G) Histograms showing the percentage of cells positive for HLA-DR activation marker expression (dark, black line) on “cytolytic” CD56^dim^CD16^high^ NK cells from 5 subjects at weeks 0, 2 and 30 of therapy with VDZ. Gates for activation marker expression was set according to FMO control staining tube which contained NK phenotypic markers, but not activation stains (shaded, gray line). Red numbers in upper right corner represent the Geometric Mean Fluorescence Intensity (GMFI) of HLA-DR in each subject.

We further looked at the expression of HLA-DR and PD-1 on CD56^dim^ cytolytic NK cells. Interestingly, pronounced increase in HLA-DR was noted in 4/5 subjects at wk 2, suggesting an activation of cytolytic NK cells post-VDZ (Figure 5E-G). Due to paucity of GI-derived cells, we were unable to assess for NK cell changes in the GI tract post-VDZ. In summary, our data demonstrates changes in NK cell frequency and increased NK cell activation in circulation, early after anti-α4β7 therapy.

### Anti-α4β7 therapy results in a variable effect on stimulated and unstimulated multiply spliced HIV-1 transcripts in blood derived CD4^+^ T cells

To study the impact of VDZ on the HIV reservoir we used a combination of approaches assessing HIV-related measurement. Importantly, all the patients continued on fully suppressive antiretroviral therapy for the course of this study. We observed a decrease in total and integrated HIV-1 DNA levels in sorted CD4^+^ T cells derived from PBMCs in 4/5 patients (Figure 6A). In the TI, while there was a decrease in total and integrated HIV-1 DNA in 3/5 subjects, in 2/5 subjects there was an increase (Figure 6B, top panels). Similarly, the effect of VDZ treatment on HIV-1 DNA levels in the LC was variable and not significant (Figure 6B, bottom panels). We did not detect HIV-1 RNA levels in unstimulated GI-derived CD4^+^ T cells and paucity of cell numbers made performance of quantitative viral outgrowth assays (QVOA) not feasible in this initial study.

**Fig. 6.**
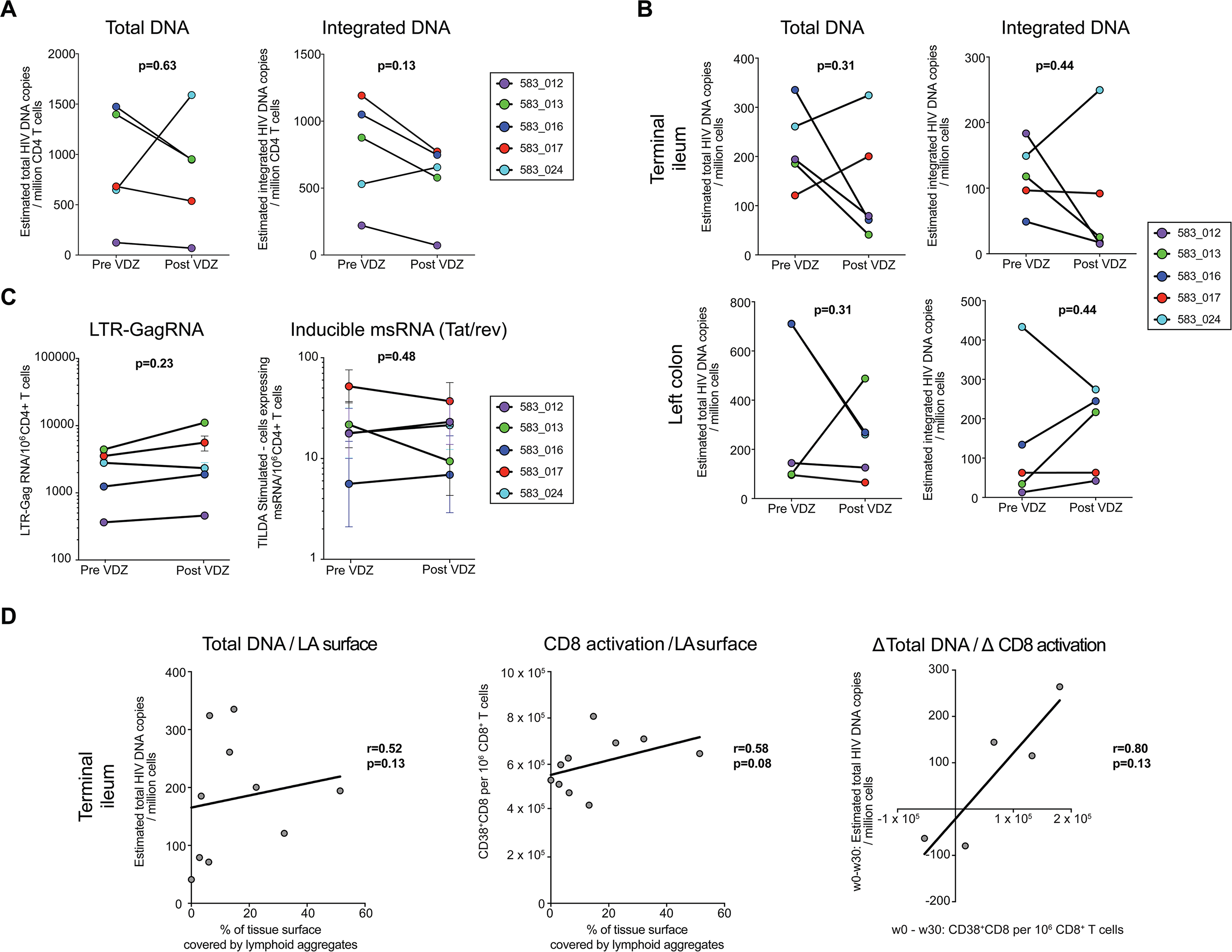
Impact of anti-α4β7 therapy on HIV-1 levels in the peripheral blood and in the GI tract. (A) Change in estimated copies per million cells of total and integrated HIV DNA in sorted CD4^+^ T cells derived from the peripheral blood. (B) Change in total and integrated HIV DNA in whole biopsies derived from the TI (top panels) and LC (bottom panels) post-VDZ. DNA levels were normalized per housekeeping gene copy number (C) Change in HIV-1 long terminal repeat (LTR)-gag RNA in unstimulated bead-selected, circulating CD4 cells pre-VDZ and at wk 30 post-VDZ (left panel). In the right panel, change in the frequency of cells with inducible multiply spliced RNA (tat/rev), measured with the TILDA assay, are compared pre- and post-VDZ. (D) Plots showing the linear regression line between various parameters in the TI mucosa. Left panel shows correlation between HIV total DNA within the TI and the % of tissue covered by lymphoid aggregates. Middle panel shows the correlation between the frequency of CD8^+^CD38^+^ activated cells and the surface covered by lymphoid aggregates in the TI. The right panel represents the correlation between the magnitude of w0 to w30 changes in total HIV-1 DNA levels and the frequency of CD8^+^CD38^+^ cells. A positive number is representative of a decrease from wk0 to wk30. Correlation factors and p-values were estimated with the Spearman correlation test.

Tat/Rev Induced limiting dilution assay (TILDA) was performed on peripheral blood derived, bead-sorted CD4^+^ T cells pre-VDZ and at week 30 post-treatment. In subjects 583-013 and 583-017, lower frequencies of cells with multiply spliced HIV RNA (tat/rev) were seen after treatment. In the absence of stimulation, Tat/rev transcripts (multiply spliced) were rare, resulting in poor reproducibility between replicates (Figure 6C left panel). Overall, in the face of ongoing cART, spliced and unspliced HIV-1 RNA om the blood showed variability and inconclusive overall effect of anti-α4β7 on markers of HIV-1 persistence (Figure 6C, right panel).

Further, we correlated the total DNA level in the TI with immunological parameters. Interestingly, we found a positive correlation (r=0.52, p-value=0.13) between the degree of lymphoid aggregates and the DNA level suggesting a potential role of the TI lymphoid structures as a viral resrvoirs. Additionally, the percentage of the GI surface covered by lymphoid aggregates in the TI correlated positively with CD8^+^ T cell activation (Figure 6D). Finally, we found a strong correlation (r=0.8) between the magnitude of CD8^+^ T cell activation and decrease in total DNA levels in the TI with post-VDZ (Figure 6D).

## DISCUSSION

A growing body of evidence has demonstrated that α4β7^+^CD4^+^ T cells represent an early target for HIV-1 (*18, 19, 21*). Accordingly, NHP studies utilizing an anti-α4β7 mAb, that is a close analogue of VDZ, either prior to infection, or in the context of SIV infected animals receiving cART, have yielded promising and provocative results (*24, 25*). This has prompted the initiation of two phase I clinical trials of VDZ in HIV infected subjects. However, the underlying mechanism(s) by which anti-α4β7 Mab therapy impacted SIV pathogenesis remains unknown. Subjects co-afflicted with HIV and IBD represent a unique opportunity to investigate the clinical utility of VDZ therapy in HIV disease. The goals of the present report, the first in human, were to determine the safety and tolerability of anti-α4β7 therapy in humans with HIV-1 infection. Additionally, we performed detailed immunological analyses in the intestinal tissue and peripheral blood in order to provide insight into the mechanism(s) of action of this drug.

Data reported herein from 6 patients with chronic HIV-1 infection (5 patients with detailed intestinal immunophenotyping) demonstrated that VDZ can be safely administered over extended periods of time. The longest duration of treatment that we have recorded in an HIV-1 infected subject is 19 months of uninterrupted a4b7 therapy without any significant adverse events. In fact, the only AEs recorded were minor and self-limited episodes of upper respiratory infection as described previously (*26*).

At the time of writing this report, we received information regarding a patient who was diagnosed with progressive multifocal leucoencephalopathy (PML) while on VDZ therapy. This patient had history of Crohn’s Disease for ∼20 years; was on treatment with azathioprine and VDZ therapy, and developed neurological symptoms that led to the diagnosis of PML. Subsequent medical work up revealed an undiagnosed HIV-1 infection with a CD4^+^ T cell count of 300 cells/mm^3^ (personal communication, Takeda Pharmaceuticals, Supplementary data).

While important details continue to emerge about this case, it is a sobering reminder of the potential for serious infectious complications in patients with HIV-1 infection. Identifying the role, if any, of VDZ in the appearance of PML in the patient above will be challenging. Advanced HIV-1 infection, which is associated with T cell dysfunction, is a known and significant risk for PML (*41*). Additionally, azathioprione, an inhibitor of T cell activation, is also a known risk-factor for PML (*42*). Looking forward, a prudent approach would be to avoid anti-α4β7 therapy in viremic HIV-1 infection, in the presence of concomitant immunosuppression in patients seropositive for John Cunningham virus (JCV).

In order to determine the immunological effects of VDZ on the GI tract, we examined both immune inductive sites (represented by lymphoid aggregates, concentrated in the TI) and effector sites (represented by lamina propria associated lymphocytes (LPLs)). Notably, inductive sites such as the Peyer’s patches harbor antigen-naïve T and B cells while effector sites such colonic lamina propria contain antigen-experienced memory T cells and plasma cells (*5*). Significantly reduced frequency of non-plasma cell B cell subsets (but not plasma cells) and naïve T cells (but not memory T cells) by flow-cytometry, post-VDZ treatment drew our attention to examining the effect of anti-α4β7 therapy on lymphoid aggregates. In confirmation of our hypothesis, we observed a striking attenuation of lymphoid aggregates in the course of treatment. These data, represent a novel mechanism of action (MOA) of anti-α4β7 therapy, and can be explained, in part, by the fact that lymphocyte homing to both inductive and effector sites, is α4β7-dependent (*6, 43*). While, the existing literature has focused on drug-therapeutic effects on memory cells homing to the effector compartment, we observed an even greater effect on the homing of naïve T and B cells to the inductive sites. Consistent with our observations, previous studies have reported variable effects anti-α4β7 on treatment on lamina propria lymphocytes (*34*). This may reflect redundant cellular homing pathways to the lamina propria and a more exclusive dependence on α4β7-MAdCAM interactions in homing to inductive sites. In support, we have observed a significant increase in β7^hi^memoryCD4^+^T cells in circulation post-VDZ therapy. Furthermore, NHP studies do not demonstrate hypocellularity in the lamina propria post- VDZ treatment which would be expected in the face of blocking cellular ingress (*44*). Viewed collectively, no significant change in CD4^+^T cell frequencies in the lamina propria concurrent with an increase in β7^hi^memoryCD4^+^T in circulation implies that CD4^+^ T cells are utilizing redundant, and perhaps α4β7-independent pathways of localizing to the GI lamina propria during VDZ therapy. A better understanding of these pathways has important implications for IBD therapeutics as well.

Attenuation of lymphoid aggregates by anti-α4β7 therapy may have important implications for HIV-1 infection. It is well established that B cell follicles are a source of viral replication in the context of chronic infection (*45*). Of note, emerging data suggests that B cell follicles are a major source of replication competent virus during cART (*46*), and may represent one barrier to viral eradication. Follicular dendritic cells (FDC), physiological long term stores of antigen for the development of high affinity antibodies (*47*) accumulate a large reservoir of infectious extracellular HIV virions within the B cell follicles and serve as a major source of infectious virions in vivo (*48-50*). Virus can then be passed to follicular CD4^+^ T cells (including TFH cells) that are highly susceptible to HIV-1 infection (*51, 52*) due to a variety of mechanisms including BCL-6 mediated diminished expression of interferon-stimulated genes (*53*). Moreover, B cell follicles may be semi-immune privileged sites due to the inability of cytotoxic T cells lacking follicular homing molecule CXCR5 (*54*) from entering them (*55*). In accordance with this concept, it has been hypothesized that overcoming the immune privilege of lymphoid follicles may be key to HIV-cure efforts (*56*). Among the strategies being considered are the development CXCR5-expressing chimeric antigen receptor (CAR) T cells (*57*), the use of bispecific antibodies (*58*) or treatment with latency reversal agents (LRA) (*59*). While significant attention has been paid to LRAs like histone deacetylase inhibitors (HDACi), protein kinase C (PKC) agonists and recombinant IL-15, to date, no LRA agent is known to specifically target the lymphoid aggregates. Here, unexpectedly we have identified that a gut-specific immunotherapeutic agent can lead to the attrition of B cell follicles in the GI tract. This represents a potentially important tool in the HIV-1 cure effort. Given that the impact of anti-α4β7 therapy is primarily GI-specific, it would possibly need to be combined with other agents to impact on HIV-1 reservoirs in both GI and non-GI sites. Another implication of the present report is the proposed duration of therapy. We contend that extended treatment with anti-α4β7 agents, either alone, or in combination with other LRAs will likely be required for a sustained impact on the lymphoid aggregates and should be considered in the design of future animal and human studies.

In addition to its effect on lymphoid aggregates, anti-α4β7 was associated with a decrease in CD4^+^ T cell activation in the terminal ileum. These data suggest an effect of immune stimulation by viral reservoirs in the terminal ileum. However, an obvious limitation here is the low number of patients- such observations would need to be validated in a larger dataset. An additional point of interest and perhaps the subject of future investigation is the impact of therapy on the activation of CD56d^im^ cytolytic NK cells at wk 2 post-treatment. Unfortunately, we do not have tissue assessments of NK cell phenotype/function to better understand the impact of therapy on this important innate immune subset.

Virologic effects of therapy, modest at best in the present study were not unexpected in the face of ongoing, fully suppressive cART. To better understand the virologic consequences, structured treatment interruption studies are being planned with detailed characterization of viral reservoirs in peripheral blood and tissue- before and after cART interruption.

The patients described herein had concomitant IBD which could potentially confound the interpretation of our results. Given that anti-α4β7 therapy is not yet licensed for clinical use in HIV alone, this was a practical limitation for this study. That said, we believe that the impact of IBD on our analysis was minimal. At the outset, the patients had very mild IBD with history of distal rectal involvement as detailed in the clinical characteristics of the cohort. Secondly, histological examination of tissues, arguably the gold standard for defining tissue inflammation confirmed that the biopsied areas within the TI or the LC did not have mucosal inflammation. Finally, non-invasive markers including CRP confirmed no significant differences between the HIV-1 infected patients and normal volunteers. Nevertheless, we are planning studies in non-IBD, HIV subjects.

In summary, the present report provides the first evidence of long-term safety and tolerability of anti-α4β7 therapy in HIV infected subjects and provides novel insights into its effect on lymphoid aggregates. This activity may represent a new approach toward impacting persistent viral replication in gut tissues, which play a central role in HIV pathogenesis. In the ongoing efforts to develop an HIV cure, we believe that treatment with anti-α4β7 agents in combination with other latency reversal agents may represent a new therapeutic approach.

## MATERIALS AND METHODS

### Study design and ethical considerations

Patients were recruited from within a cohort of HIV-1 infected patients with concomitant IBD (*29*) being followed at the Icahn School of Medicine at Mount Sinai and its affiliated hospitals and clinics. Informed consent was obtained from all the participants. The study protocol was approved by the Mount Sinai institutional review board.

6 subjects with IBD and concomitant HIV-1 infection were prospectively enrolled into the study. Of these, ileocolonoscopy was performed pre-VDZ initiation in 5 subjects. In one subject, colonoscopy could not be performed pre-treatment. Therefore, full immunological and virologic analyses are being reported on 5 subjects. VDZ infusions were administered following the FDA approved protocol for IBD, including an induction phase with a 300 mg iv infusion at week 0 (wk0), wk2, wk6 subsequently followed by a maintenance phase with a 300 mg iv infusion every 8 weeks. Before each infusion, blood was collected for analysis. A repeat colonoscopy was performed on each of the 5 subjects between wk 22 and wk 30 post-treatment. While the patients continue to be on long-term follow-up and on VDZ therapy, for the present report, we are describing the effects of 30 weeks of VDZ therapy.

In addition, we recruited a cohort of healthy volunteers (n=12) (clinical details in Table S1). All healthy volunteers and Chronic HIV-1 subjects underwent colonoscopy and phlebotomy for immunological analyses in the GI tract and peripheral blood respectively. Among the 6 patients, TI and LC biopsies were obtained both pre and post VDZ initiation for 5 of them. All the patients continued stable and uninterrupted cART for the length of the study. Importantly, at the time of writing this manuscript, the patients continued to be on cART as well as VDZ.

### Peripheral blood mononuclear cell (PBMC) isolation

50 ml of blood were drawn in EDTA tubes at baseline and preceding each VDZ infusion. Blood was processed, under sterile condition, within 2 hours after collection. Plasma was collected and stored immediately at −80°C. PBMCs were obtained using standard Ficoll Hypaque centrifugation gradient. Briefly, whole blood was diluted 1:1 with phosphate buffered Saline (PBS, Gibco) and gently overlaid on lymphocyte separation medium^®^ (MP Biomedical). After centrifugation, the phase containing PBMCs was harvested and washed 2 times with sterile PBS. A subset of the PBMCs was directly used for staining while the remaining was cryopreserved at - 150C using standard techniques.

### Isolation of Gastrointestinal mononuclear cells

During the ileocolonoscopy procedure, mucosal biopsies were obtained using standard large cup biopsy forceps. Notably, we harvested biopsies from the TI as well as the uninflamed LC (n=30, limiting the effect of variability in sampling). Fresh biopsies were transported in a medium containing RPMI (GIBCO) and processed immediately using standard techniques in the lab (*2*). Briefly, the intestinal biopsies were transferred to 10 ml of RPMI containing 0.005 mg of collagenase (Sigma-Aldrich) and incubated for 22 minutes with gentle agitation (215 rpm) at 37°C. Biopsies were then physically disrupted by chopping with scissors until obtaining a “porridge-like” consistency. The chopped tissue was transferred into a fresh RPMI-collagenase solution for a second round of 22-minute incubation at 37°C with gentle agitation. After incubation, the single-cell suspension was obtained by filtering through a 100-µm and 40- µm cell strainer and washed with RPMI twice.

### Flow cytometry analysis

Isolated cells were incubated with respective live/dead dye and antibodies for 20 min at 4°C in the dark, washed 2 times with PBS and then fixed using Cytofix/Cytoperm buffer (BD) until acquisition. Samples were acquired using LSR Fortessa (BD), and data were analyzed using FlowJo v10 (FlowJo). Dead cells and doublets were excluded from all analyses. The list of used Abs is displayed in Supplementary table 2.

### Immunohistochemistry

Biopsies were collected directly in formalin and fixed for 24-48 hours. Fixed tissue was then embedded in paraffin. Tissue sections were immunohistochemically (IHC) stained with CD20 and CD4 antibodies. 4 µm paraffin sections were deparaffinized, and this was followed by antigen retrieval in EDTA buffer (pH 9.0) (Er2, Leica) for CD4 and in low pH (pH 6.0) citrate solution (Er1, Leica) for CD20. The rest of the procedure was done in a Leica bond 3 automated instrument for CD20 and CD4 single staining. Endogenous peroxidase activity was blocked with 3% hydrogen peroxide. The slides were incubated with pre-diluted antibodies against CD20 (clone L26, Bond RTU Primary, Leica) or CD4 (clone 4B12, Bond RTU Primary, Leica) at room temperature for 1 hour. The detection was done using the Bond™ PolymerRefine Detection system (Leica) including a hematoxylin counter stain. Frequency of CD20 and CD4 within the lamina propria was then counted in a blinded fashion by HK and MT and expressed as positive cells per high power field.

CD19 - CD4 double staining was performed using the Ventana Discovery ULTRA platform. Antigen retrieval was done at a slightly basic pH with CC1 reagent (Ventana). A mouse anti-human CD4 (clone 4B12, Agilent) and a rabbit anti-human CD19 (clone EPR5906, abcam) were used as primary antibodies. For DAB, anti-mouse HRP pre-diluted secondary antibody (Ventana) was used and then Discovery ChromoMap DAB was applied (Ventana). For purple stain, we used anti-Rabbit HQ (Ventana) secondary, then applied anti-HQ HRP (Ventana) followed by the application of Discovery purple color (Ventana).

### Immunofluorescence assay

Tissues were fixed in 10% neutral buffered formalin, and immunofluorescence was performed on 5µm paraffin-embedded section. Deparaffinization and antigen retrieval were performed by heating sections 55°C for 20 minutes, followed by a series of hydration washes as follows: three 100% Xylene for 5 minutes each, two 90% EtOH for 3 minutes each, 70% EtOH for 3 minutes, 30% EtOH for 3 minutes, diH_2_0 for 3 minutes. Antigen retrieval was achieved by boiling sections in citrate buffer (pH=6.5) for 10 minutes. Tris Buffered Saline (TBS) supplemented with 0.05% Tween 20 was used as a wash solution and TBS supplemented with 0.1% Tween 20 with 2.5% bovine serum albumin (BSA) was used as an antibody diluent. Primary antibodies for CD20 (polyclonal Goat Ab, Sigma Aldrich) and CD3 (clone SP7, Rabbit mAb, Abcam) were diluted with diluent to 1:200 and 1:100 respectively and incubated at room temperature for 1 hour. Secondary antibodies for CD20 (Alexafluor 488 Donkey anti Goat 705) and CD3 (Alexafluor 594 Donkey anti Rabbit) were diluted with diluent to 1:200 each and incubated at room temperature for 1 hour. Slides were mounted with Vectashield Hardset with DAPI and allowed to cure for 1 hour before storing slides at 4°C. Slides were analyzed using an AxioImager Z2 microscope (Zeiss) and Zen 2012 software (Zeiss).

### Cell-associated HIV viral DNA measurements

*For the peripheral blood derived CD4 T cell DNA/RNA isolation* cryopreserved PBMCs were rapidly thawed and enriched for CD4^+^ T cells to high purities with an EasySep Human CD4^+^ T cell enrichment kit (Stemcell Technologies). Both DNA and RNA were extracted from enriched CD4 T cells using AllPrep DNA/RNA/miRNA Universal Kit (Qiagen). DNA and RNA purity and quantity were measured using the ND-2000 spectrophotometer (NanoDrop Technologies).

*For the gastrointestinal tissue (Ileal and sigmoid colonic) DNA/RNA isolation,* 4 to 6 biopsies were placed in RNALater (Ambion) and stored at −80°C until nucleic acid isolation. Tissue biopsies were homogenized using the Qiagen TissueRuptor II in Qiagen RLT Buffer. DNA and RNA were isolated from homogenized tissue biopsies using the Qiagen AllPrep DNA/RNA/miRNA Universal kit according to manufacturer’s protocol for tissue specimens. *Determination of total cellular and integrated HIV-1 DNA* was conducted according to previously published methods (*60*). In brief, pre-amplification using LTR-gag-specific primers ULF1 (5’ ATG CCA CGT AAG CGA AAC TCT GGG TCT CTC TDG TTA GAC 3’; HXB2 452-471) and UR1 (5’ CCA TCT CTC TCC TTC TAG C 3’; HXB2 775-793) for total HIV-1 DNA, and using LTR-Alu specific primers ULF1, Alu1 (5’ TCC CAG CTA CTG GGG AGG CTG AGG 3’) and Alu2 (5’ GCC TCC CAA AGT GCT GGG ATT ACA G 3’) for integrated HIV-1 DNA was conducted. Pre-amplification process was carried out using a reaction volume of 50 μl with 5 μl Taq Buffer, 3 μl of MgCl_2_, 0.6 μl of each dNTP, 1.5 μl of each primer, 0.5 μl of Taq polymerase and 5 μl of DNA template. Cycling conditions are 95°C for 8 min, then 12 cycles of 95°C for 1 min, 55°C for 40s (total DNA) and 1 min (integrated DNA) and 72°C for 1 min (total DNA) and 10 min (integrated DNA), followed by elongation at 72°C for 15 min. The pre-amplification process was performed on a GeneAmp PCR System 9700 (Applied Biosystems Inc). For the subsequent total and integrated HIV-1 DNA quantification, samples were quantified with a qPCR TaqMan assay using Lambda T (5’ ATG CCA CGT AAG CGA AAC T 3’) and UR2 (5’ CTG AGG GAT CTC TAG TTA CC 3’; HXB2 583-602) coupled with a FAM-BHQ probe (5’ CAC TCA AGG CAA GCT TTA TTG AGG 3) on a StepOne Plus Real-time PCR System (Applied Biosystems Inc). Cell-associated total and HIV DNA quantification was measured using a reaction volume of 20 μl with 10 μl of 2x TaqMan Universal Master Mix II including UNG (Life technologies), 0.25 μl of 100 μM each primer, 0.4 μl of 10 μM probe, and 5 μl of 1:10 diluted pre-amplified PCR product as template. Cycling conditions were 50°C for 2 min, 95°C for 10 min, then 60 cycles of 95°C for 15s and 60°C for 1 min. Cells-equivalent DNA copy number was determined using CD3 gene, and CD3 DNA copy number was quantified by direct extrapolation against a standard curve.

*For the quantification of cell associated total HIV-1 RNA,* qPCR TaqMan assay using LTR-specific primers F522-43 (5’ GCC TCA ATA AAG CTT GCC TTG A 3’; HXB2 522–543) and R626-43 (5’ GGG CGC CAC TGC TAG AGA 3’; 626–643) coupled with a FAM-BQ probe (5’ CCA GAG TCA CAC AAC AGA CGG GCA CA 3) on a StepOne Plus Real-time PCR System (Applied Biosystems Inc) was conducted. Cell associated HIV-1 RNA copy number was determined in a reaction volume of 20 μl with 10 μl of 2x TaqMan RNA to Ct 1 Step kit (Life Technologies), 4 pmol of each primer, 4pmol of probe, 0.5 μl reverse transcriptase, and 5 μl of RNA under identical cycling conditions. Cells-equivalent RNA copy number will be determined RPLP0 gene. Patient specimens were assayed with up to 800 ng total cellular RNA or DNA in replicate reaction wells and copy number determined by extrapolation against a 7-point standard curve (1–10,000 cps) performed in triplicate.

### Quantification of cell-associated LTR-Gag RNA

Total CD4 T cells were isolated from frozen PBMCs using negative magnetic selection (Stem Cell technologies). RNA was extracted from isolated CD4 T cells with the Qiagen RNeasy Mini kit adding the Dnase step. Cell-associated LTR-gag transcripts, were measured by nested real-time PCR. The primers used were previously described in Vandergeeten et al. (*60*). Results were expressed as HIV RNA copy numbers per million CD4^+^ T cells.

### Tat/Rev induced Limiting dilution Assay (TILDA)

The frequency of CD4+ T cells expressing multiply-spliced HIV RNA upon stimulation was measured using the Tat/Rev Induced Limiting Dilution Assay (TILDA) as previously described (*61*). Briefly, CD4 T cells were isolated from cryopreserved PBMCs using negative magnetic selection (Stem Cell technologies) and stimulated with PMA (100ng/mL) and ionomycin (1µg/mL) for 18 hours. Stimulated cells were serially diluted and the wells in which cells producing tat/rev RNA were identified by semi-nested real-time PCR.

### Statistical analysis

Plots were drawn using GraphPad Prism software^®^. Statistical significance of immunophenotyping and viral data was assessed using the two-sample paired Wilcoxon signed rank test and 2-tailed (paired) student t test when appropriate. Correlations were assessed using the Spearman test.

## SUPPLEMENTARY MATERIALS

Fig. S1. All sites sampled for immunological and virological analyses were histologically uninflamed.

Fig. S2. B cell gating strategy.

Fig. S3. CD20 immunostaining of terminal ileum biopsies of all patients, pre and post VDZ.

Fig. S4. T cell gating strategy

Fig. S5. NK cell gating strategy

Table S1. Clinical characteristics of healthy volunteers.

Table S2. Flow cytometry antibodies.

## ACKNOWLEDGEMENTS

We thank the patients who participated in the study. The authors would like to thank the Biorepository and Pathology Core at Icahn School of Medicine at Mount Sinai for carrying out some of the immunostaining experiments. **Funding**: This work was supported by the following grants: NIH/NIDDK R01 112296 (SM), R01 DA041765 (SM, BKC, ALP) and NIH/NIAID grant UM1 AI126620 (LJM). Additional support was provided by The Philadelphia Foundation (Robert I. Jacobs Fund), Kean Family Professorship, the Penn Center for AIDS Research (P30 AI 045008). MU has received fellowships from la Fondation pour la Recherche Médicale (FDM 41552) and from la Société Nationale Française de Gastro-Entérologie (SNFGE). AKR was supported by the Digestive Disease Research Foundation (DDRF). **Author contributions**: MU, AKR, MT, LL, CT, IS, MC, EP, AC, ALP and ID designed and carried out experiments. HMK, HU, IDI, BKC, TWS, JKL, LJM, NC, LCN and SM supervised experiments, analyzed and interpreted data. EKL, GR, JA, IO, JFC and SM recruited study subjects and provided samples. MU and SM drafted the manuscript. SM designed the study, supervised experimental data collection and coordinated integration of collaboration between all participating laboratories. All authors critically reviewed and edited the final version of the manuscript. **Competing interests**: SM and JFC have an unrestricted, investigator-initiated grant from Takeda Pharmaceuticals to examine novel homing mechanisms to the GI tract.

